# Critical Variables to be Considered when Attempting to Estimate Blood Ethanol Concentrations in Rats from g/kg Exposure Data

**DOI:** 10.1101/582361

**Authors:** Carol A. Dannenhoffer, Dominika Hosová, Sanjeena Dang, Utkarsh Dang, Linda P. Spear, Neurobiology of Adolescent Drinking in Adulthood (NADIA) Consortium

**Author notes:** Communicating Author: Carol A. Dannenhoffer Bowles Center for Alcohol Studies UNC Chapel Hill, NC. The work presented in this manuscript was funded by grant AA017823 to Linda P. Spear and T32 Training Grant 1T32 AA025606-01 at Binghamton University.

## Abstract

**Background:** In rodent studies of ethanol (EtOH) consumption where blood sample collection does not occur, there is often mention of likely BECs based on prior studies. These studies may vary in dose(s) used, age/sex/species, or administration route. Often, intake studies may presume that binge-levels were achieved without knowing that BECs exceeded 80 mg% (binge threshold). In human studies, estimated BECs (eBECs) have been derived using complex formulas that consider EtOH consumption level and the weight and sex of the individual.

**Method:** Three datasets were used to derive eBECs using a conversion factor (CF) that considers gram (g) of EtOH per kilogram (kg) of animal weight and other variables that may influence BECs such as age, sex, dose, route, vehicle, chronicity, and timing post-exposure. Regression analyses were also conducted for each dataset, building regression models with BEC as the response and other variables in the study specific to each dataset as predictor variables.

**Results:** Dataset1 assessed age, sex and post-injection time point. Both CF and regression analyses determined that different CFs should be used for 10- and 30-min post-administration time points. Dataset2 assessed age, dose, vehicle and post-intubation time point. Depending on the post-intubation time point, several CFs were used to derive eBECs. When weight was not used as a regression variable, data across approaches corresponded, with age differences emerging later in elimination phase. In Dataset3 that used BECs from a repeated intake study, chronic exposure influenced CFs, although regression analysis did not yield similar findings.

**Conclusions:** Although eBECs can be derived, critical variables vary with subject and test conditions and do not always concur with results of regression analyses. Although, not designed to replace assessment of BECs when sample collection is possible, the CF approach may prove useful when estimating BECs in studies where assessments are not feasible.

## Introduction

When interpreting data generated in alcohol studies, it is often desirable to anchor the findings in blood ethanol (EtOH) concentrations (BECs). This is especially the case when relating relative exposure levels to “binge” consumption levels defined by NIAAA in terms of the BECs produced (≥ 80 mg %). Although preferably BECs should be determined in each study, sometimes such assessments are not feasible, and estimated BECs are utilized. In human studies, estimated BECs are typically derived using a conversion-factor-based formula based on the consumed volume, body weight, and sex (Pizon et al., 2007). In rodent studies, when bloods are not collected for BEC determinations, BECs are often roughly estimated by reporting BEC data from other studies observed at seemingly comparable EtOH exposure levels. However, estimates obtained across studies in laboratory animals may not consider critical variables (e.g., administration route, age, sex, etc.) that could potentially influence BECs.

The present study assessed the degree to which BECs can be reliably estimated across a variety of such variables. Three different datasets from our lab were analyzed to explore effects of variables such as age, sex, EtOH vehicle, chronicity, route and post-exposure-to-assessment timing when attempting to estimate BECs. Age and sex may be important variables for consideration when deriving eBECs given that these variables are known to influence pharmacokinetics and body composition (Marshall et al., 1983, Vestal et al., 1977). Indeed, sex and the time over which alcohol was consumed are the only variables included in NIAAA’s estimate of the number of drinks required to reach the binge drinking threshold, with the > 80 mg% cut-off for binge drinking stated as being typically reached in humans after consumption of 5 standard drinks in males and 4 in females within a 2 hour period. Age may also play an important role, with greater rates of EtOH metabolism but lower percentages of body water (within which EtOH distributes) in youth than in adults (see (Spear, 2007, Donovan, 2009) for discussion). Indeed, based on these considerations, Donovan (2009) recommended that binge drinking among youth be defined as 3 or more drinks in 2 hours for 9-17 year old girls, and 3, 4 or 5 drinks within a 2 hr period for 9-13, 14-15 or 15-17 year old boys, respectively. These sex and age differences in threshold number of drinks are in part related to body weight differences, with females generally being lower in weight than males, and body weights increasing across the adolescent period into adulthood. Body weight differences are controlled in rodent studies via routinely expressing EtOH exposures in g/kg body weight. Nevertheless, age and sex differences in EtOH metabolism/elimination have also been reported in rodent studies, with adolescent rats sometimes (Hollstedt et al., 1977, Brasser and Spear, 2002) but not always (Kelly et al., 1987, Silveri and Spear, 2000) reported to exhibit modestly but significantly higher rates of metabolism of EtOH and other drugs than adults (see (Spear, 2007) for review and references). Elimination of EtOH has also been reported to be about 15% more rapid in female than male rats (Robinson et al., 2002), although differences in BECs after EtOH challenge to males and females are not ubiquitous (e.g., (Goist Jr and Sutker, 1985)).

When estimating BECs, it is also important to consider the pharmacokinetics of EtOH. The timing of peak EtOH concentrations and EtOH clearance are dependent upon many factors, including dose (Wilkinson et al., 1977), chronicity (Winek and Murphy, 1984) and route of administration (Wilkinson, 1980). Repeated exposure to alcohol induces various types of tolerance including both functional (Broadwater et al., 2011) and metabolic (Silvers et al., 2003) tolerance. To the extent that metabolic tolerance develops, BECs would be expected to be lower following EtOH challenge in animals with a history of EtOH exposure relative to animals without prior exposure (e.g., (Silveri and Spear, 2001)). When EtOH first enters the body, there is a concentration gradient that is highest at the route of entry and spreads across the water compartment of the body; as this gradient reaches equilibrium, the flow inverts as EtOH is gradually eliminated (Norberg et al., 2003). Different routes of administration induce different time to peak levels and different concentration gradients; for instance, the gradient caused by the absorption of digesting EtOH from the stomach after oral consumption or intragastric (i.g.) intubation is subject to first past metabolism (Julkunen et al., 1985) and is slower, with longer times to peak, than when EtOH is injected directly into the intraperitoneal (i.p.) cavity (Spirduso et al., 1989). Hence, the peak BEC reached, as well as the time to peak and rate of elimination, likely varies across route of exposure. Whether EtOH is self-administered or administered as a bolus via gavage is also likely to influence peak BEC levels (see (Hosová and Spear, 2017) for discussion). The vehicle used may also matter. Our lab uses both supersaccharin (“supersac”: 3% sucrose + 0.125% saccharin in water (Ji et al., 2008)) and chocolate Boost^®^ to increase voluntary consumption in EtOH intake studies. While both are palatable (Hosová and Spear, 2017), the chocolate Boost^®^ solution contains more nutrients relative to supersac (in 100 ml of 25% EtOH: [Supersac = 8.89 calories; chocolate Boost^®^ = 74.63 calories]), thus potentially altering the pharmacokinetics of EtOH (Cederbaum, 2012), a possibility that was also examined.

In the present study, two different approaches were used to derive converging evidence for the impact of the various independent variables on BECs in the rat. The first strategy derived conversion factors (CFs) to estimate BECs (eBECs) via dividing each animal’s BEC by the amount of EtOH administered or ingested, and using the average of these CFs to derive eBECs that were compared with actual BECs, with additional variables considered as necessary to improve the fit. The second strategy used a modeling approach to build regression models with BEC as the response and other variables in the study (specific to each dataset) as predictor variables.

## Materials & Methods

### General Method

#### Subjects

Sprague Dawley rats used to derive these datasets were obtained from our colony at Binghamton University. All rats were culled to 8-10 pups per litter one day after birth (postnatal day [P] 1) and were housed with one or more same-sex littermates on P21. All animals received Purina Rat Chow (Lowell, MA) and water ad libitum and were maintained on a 12-/12-hr light/dark cycle (lights on at 0700h). The subjects were handled throughout the experiments under guidelines established by the National Institute of Health, using protocols approved by the Binghamton University Institution Animal Care and Use Committee.

#### Dataset Recruitment

Three datasets from our lab were used in the present investigations:

Dataset1 was from a study we conducted to explore sex and age effects on BECs at different lengths of time (10 and 30 minutes) following an i.p. injection. Both male and female adolescent (P35) and adult (P70) rats were injected i.p. with a moderate dose of EtOH (1 g/kg) and trunk blood was collected for BEC analysis 10 or 30 minutes later. These data were used to explore the impact of age, sex and post-injection interval on eBEC determination.

Dataset2 was derived from a study that examined BECs after administering EtOH solutions varying in caloric content (using either chocolate Boost^®^ or supersac as the vehicle). EtOH was administered via i.g. to adolescent (P33-36) and adult (P75-84) males, and trunk blood collected for BEC assessment at three different time points. Dataset2 was included in this analysis to examine the contribution of age, vehicle and timing on the determination of eBECs.

Dataset3 used a model of binge-level voluntary drinking we developed in adolescent Sprague-Dawley rats (Hosová and Spear, 2017), with the dataset chosen for analysis from a follow-up study comparing intake and BECs in male and female adolescent rats when using ball-bearing-containing versus open-ended sipper tubes (sample sizes of 8/group). For a total of 14 access days (P28-41), half of the adolescent rats in the study were given 30 minutes of access to Boost^®^ solution from bottles containing open-ended sipper tubes while access for the other half was provided by ball-bearing-containing sipper tubes. Following the drinking session, blood samples for BEC analysis were collected from the lateral tail vein of each animal on two separate occasions (after days 6 (P33) and 14 (P41)). This dataset was used to address the impact of sex and age/chronicity on eBECs after voluntary consumption in adolescent rats.

#### Analysis of BECs

Depending on the experiment, trunk or tail vein blood samples were collected either following rapid decapitation or after lateral tail vein incision, respectively. Whole blood samples were stored at −80°C until time of BEC analysis via headspace gas chromatography using a Hewlett Packard (HP) 5890 series II Gas Chromatograph (Wilmington, DE) and procedures in routine use in our laboratory (e.g., see (Broadwater et al., 2011)). At the time of assay, blood samples were thawed and 25 μl aliquots were placed in airtight vials, which were then placed in an HP 7694E Auto Sampler that heated each vial for 8 minutes prior to extracting and injecting a 1.0 ml sample of the gas headspace into the gas chromatograph. EtOH concentrations in each sample were determined using HP Chemstation software, which compares the peak area under the curve in each sample with those of standard curves derived from reference standard solutions.

#### CFs and eBECs

Though not the primary focus of this study, group differences in BECs were first assessed via analysis of variance (ANOVA) to assess effects of the examined variables on actual BEC levels. We then turned to calculating eBECs for comparison with actual BEC levels. First, each animal’s BEC was divided by delivered dose or amount ingested (g/kg), with the average of the resulting values used as a CF for each entire dataset. This overall CF was then multiplied by each animal’s actual g/kg EtOH exposure to derive an eBEC for each animal. Unless otherwise noted, eBECs were compared with actual BECs using repeated measures ANOVAs and the full factorial design for each study. A lack of main or interactive effects involving the variable comparing BEC to eBEC in these overall ANOVAs was interpreted as support for a single CF for the dataset. In contrast, if the ANOVA revealed significant effects involving interaction(s) of the BEC/eBEC similarity variable and one or other variables, separate CFs were calculated as necessary for each group, collapsing across non-significant variables (e.g., age or sex) where possible. Table 1 provides a summary of all CFs per dataset and the eBECs that corresponded.

**Table 1.** Summary of all Datasets: Average BECs and eBECs collapsed across non-influential conditions

| <u>Dataset</u> | <u>Route</u> | <u>Age</u> | <u>Dose</u> | <u>Time point</u> | <u>CF</u> | <u>Average BEC</u> | <u>Average eBEC</u> |
| --- | --- | --- | --- | --- | --- | --- | --- |
| 1 | i.p. | P35, P70 | 1 g/kg | 10 min | 120 | 119.5 ± 1.9 | 120 |
| 1 | i.p. | P35, P70 | 1 g/kg | 30 min | 96 | 95.6 ± 2.8 | 96 |
| 2 | i.g. | P34-36<br>P75-85 | 2 g/kg | 45 min | 57 | 114.5 ± 5.3 | 114 |
| 2 | i.g. | P34-36<br>P75-85 | 4 g/kg | 45 min | 45 | 179.5 ± 10.6 | 180 |
| 2 | i.g. | P34-36<br>P75-85 | 2 g/kg<br>4 g/kg | 90 min | 37 | 102.9 ± 5.1 | 101.5 ± 3.9 |
| 2 | i.g. | P34-36 | 2 g/kg | 180 min | 9 | 17.1 ± 6.4 | 18 |
| 2 | i.g. | P34-36 | 4 g/kg | 180 min | 26 | 104.4 ± 14.6 | 104 |
| 2 | i.g. | P75-85 | 2 g/kg | 180 min | 23 | 45.8 ± 5.2 | 46 |
| 2 | i.g. | P75-85 | 4 g/kg | 180 min | 31 | 125.1 ± 11.3 | 124 |
| 3 | Voluntary intake | Day 6 (P33) | ~ 4 g/kg | ~30-35 min from start of drinking | 20 | 84.2 ± 8.6 | 83.9 ± 5.1 |
| 3 | Voluntary intake | Day 14 (P41) | ~ 4 g/kg | ~30-35 min from start of drinking | 14 | 61.9 ± 6.2 | 57.9 ± 3.1 |

#### Regression analyses

For this approach, we built an initial regression model with BEC as the response and other variables in the study (specific to each dataset) as predictor variables. Based off of the initial model, model assumptions were evaluated and some data points were identified as outliers and removed if necessary. When the normality assumption was not satisfied, a Box-Cox transformation (Box and Cox, 1964) on the response variable was performed to identify an appropriate transformation of the response variable to help satisfy the normality assumption of the errors. Then, using the transformed BEC as the response and other variables as predictor variables, multivariate regression analysis was performed. To identify variables that had a significant effect on the BEC values, a Bayesian information criterion (BIC; (Schwarz, 1978)) based hybrid (both) forward and backward directional stepwise variable selection approach was used. Regression equations along with summaries of results are provided. The analyses were conducted using the statistical software R (R Core Team, 2016).

### Dataset1 - CFs and BECs following i.p. administration

#### Design

The experiment was a 2 age (adolescent [P35] or adult [P70]) by 2 sex (male or female) by 2 post-injection (10 or 30 minutes) factorial design, with 6 animals per group.

##### Procedure

Animals were weighed and injected with 1 g/kg of 12.6% EtOH in 0.9% saline (v/v) i.p. and placed into their home cage until time of blood collection. At either 10 or 30 minutes post-injection, animals were rapidly decapitated and trunk blood was collected and stored at −80°C for future analysis.

#### Dataset2 – CFs and estimated BECs after i.g. intubation

##### Design

The design of this experiment was a 2 age (adolescent [P34-36] or adult [P75-85]) by 2 solution (chocolate Boost^®^ or supersac) by 2 dose (2 g/kg or 4 g/kg) by 3 time points (45, 90, or 180 minutes post-intubation) factorial with 8-10 male animals per group.

##### Procedure

To assess whether these frequently used vehicles in our laboratory for ingestion studies produced comparable BECs and CFs, in this study a 25% v/v EtOH solution was prepared in a vehicle of either chocolate Boost^®^ or supersac. Animals were weighed and intubated with 2 or 4 g/kg of EtOH in one of the two vehicles and held in a quiet room in their home cage until sample collection. Following one of the three time points post-intubation, animals were rapidly decapitated and trunk blood was collected and stored for BEC analysis.

#### Dataset3 – CFs and estimated BECs after voluntary ingestion

##### Design (as described in Hosová & Spear, 2017)

The design of this study was a 2 sex (male or female) X 2 tip-type (ball-bearing bottle tips versus open-ended tips) X 2 day (P33 or P41) repeated measures design, with 8 animals assigned to each of the 4 groups defined by the sex X tip-type factorial design.

##### Procedure (as described in Hosová & Spear, 2017)

Pair-housed animals were separated in their home cage via a wire mesh divider. Animals were given voluntary access to 10% EtOH in chocolate Boost^®^ solution (v/v) for a 30 minute period every day for 14 days beginning at P28 and ending on P41. Tail bloods were taken on days 6 and 14 (P33 and P41, respectively). Animals in this study were given the test solution in tubes containing either ball-bearing (BB) tips or open-ended (OE) tips, and were acclimatized to these tip types via receiving water in their home cages with the same type tip beginning at the time of weaning.

## Results

### Dataset1 – CFs and BECs following i.p. administration

#### BECs

Average BECs for each group are provided in Supplementary Table 1. The overall ANOVA on BECs yielded only a main effect of time point (F (1, 40) =49.928; p<0.001) suggesting that BECs were significantly higher at the 10 min (119.5 ± 1.9 mg/dl) relative to the 30 min (95.6 ± 2.8 mg/dl) time point (see Supplemental Table 1).

#### CFs and eBECs

The average of all individual CFs of Dataset1 animals was 108. Due to only one dose (1 g/kg) being administered in this study, there was no variance in eBEC because eBEC is calculated as CF x dose; therefore, every eBEC was equal to 108. The lack of variability for this variable precluded use of a repeated measure ANOVA. Instead, a difference score between BEC and eBEC was calculated for each animal and the resultant scores analyzed via a factorial 2 sex x 2 age x 2 time point ANOVA. This analysis revealed a main effect of time point (F (1, 40) =49.928; p<0.001), suggesting that the differences between estimated and actual BECs varied with the injection-to-collection time interval. Consequently, two separate CFs were calculated for these animals, one for each time point (10 min = 120; 30 min = 96) and eBECs were recalculated for each animal using the CF at the appropriate time point. No significant main effects or interactions involving the difference between estimated and actual BECs emerged at either time point, suggesting that, regardless of age and sex, these CFs were sufficient for generating appropriate eBECs at both time points after i.p. injection (10 min: BEC = 119.5 ± 1.9 mg/dl; eBEC = 120 mg/dl; 30 minutes: BEC = 95.6 ± 2.8 mg/dl; eBEC = 96 mg/dl).

#### Results of regression analysis

An initial regression model was build using age, sex and time point as the predictor variable and BEC values as the response variable. Indicator variables were created for categorical variables sex (sex = 0 for females and 1 for males) and time point (time point = 0 for 10 minutes and 1 for 30 minutes). Using a Box-Cox transformation on the response, a square transformation for the BEC values was done. Variable selection using BIC-based stepwise approach identified only the time point as a significant predictor variable at the 5% significance level (see Table 2).

**Table 2.** Regression model for Dataset1 consisting of influential conditions including time point.

|  | <u>Coefficient</u> | <u>Standard Error</u> | <u>t-value</u> | <u>p-value</u> |
| --- | --- | --- | --- | --- |
| <u>Intercept</u> | 13749.8 | 381.5 | 36.040 | < 2e-16 |
| <u>Time point</u> | -4712.7 | 533.6 | -8.831 | 3.26e-11 |

According to Table 2, the final model is:

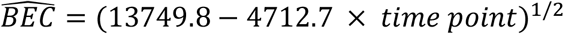

with an adjusted R-square of 0.64. Based on this regression model, for example, the estimated expected BEC at 10 minutes is 117.26 mg/dl and at 30 minutes is 95.06 mg/dl.

### Dataset2 – CFs and estimated BECs after i.g. intubation

#### BECs

Average BECs for each group are provided in Supplementary Table 2. The omnibus ANOVA of these BEC data revealed only significant main effects of dose (F (1, 179) =141.354; p<0.001) and time point (F (2, 179) =49.832; p<0.001). BECs, of course, were significantly higher at 4 g/kg (147.7 ± 6.1 mg/dl) than at 2 g/kg (75.2 ± 4.4 mg/dl). All three time points significantly differed from one another in a step-wise pattern with 45 minutes yielding the highest BECs (45 min: 147.0 ± 7.0 mg/dl; 90 min: 111.3 ± 6.4 mg/dl; 180 min: 73.9 ± 7.3 mg/dl) (see Supplemental Table 2).

#### CFs and eBECs

Averaging across all animals in this dataset, one CF (37) was calculated and used to estimate BECs. A repeated measures ANOVA comparing BEC to eBEC revealed a significant effect of time point (F (2, 179) =49.832; p<0.001), suggesting that each time point should be investigated separately with its own CF. At the 45 minute time point, using a single CF (51) led to an interaction between BEC versus eBEC and dose; therefore, different CFs for 2 g/kg (CF = 57) and 4 g/kg (CF = 45) were used. Analysis of the data using eBECs derived from these 2 CFs yielded no significant differences between BEC and eBEC (2 g/kg: BEC = 114.5 ± 5.3 mg/dl; eBEC = 114 mg/dl; 4 g/kg: BEC = 179.5 ± 10.6 mg/dl; eBEC = 180 mg/dl). At the 90 minute time point, the use of a single CF of 37 produced no significant differences between BEC and eBEC (BEC = 102.9 ± 5.1 mg/dl; eBEC = 101.5 ± 3.9 mg/dl). At the 180 minute time point, one CF (22) yielded an interaction of BEC type (estimated or actual) and dose, as well as BEC type and age; therefore, different CFs were calculated for adolescents given 2 g/kg (CF = 9), adolescents given 4 g/kg (CF = 26), adults given 2 g/kg (CF = 23) and adults given 4 g/kg (CF = 31). When each of these CFs was used to calculate eBECs for animals within each of these groups and the data analyzed as before, no significant differences between actual BECs and eBECs emerged in any of the groups (adolescents at 2 g/kg: BEC = 17.1 ± 6.4 mg/dl; eBEC = 18 mg/dl; adolescents at 4 g/kg: BEC = 104.4 ± 14.6 mg/dl; eBEC = 104 mg/dl; adults at 2 g/kg: BEC = 45.8 ± 5.2 mg/dl; eBEC = 46 mg/dl; adults at 4 g/kg: BEC = 125.1 ± 11.3 mg/dl; eBEC = 124 mg/dl). Thus, near the peak in BECs (45 min), an accurate eBEC only required a CF that was specific to each dose; at the 90 min time point, even this was no longer necessary and a single CF sufficed to estimate BECs regardless of age and dose. Later in the elimination phase (180 min), an age effect was evident, with both dose and age factors being necessary for accurate estimation of BECs.

#### Results of regression analyses

##### Method 1

An initial regression model was built using age, vehicle type, dose, weight, and time point post-intubation as the predictor variable and BEC values as the response variable. Indicator variables were created for categorical variables (age = 0 for adolescent and 1 for adult, vehicle = 0 for Boost^®^ and 1 for supersac, dose = 0 for 2 g/kg and 1 for 4g/kg, time_90 = 1 for 90 minutes and 0 for 45 and 180 minutes, and time_180 = 1 for 180 minutes and 0 for 45 and 90 minutes). We also examined two-way interactions among the predictor variables. Using a Box-Cox transformation on the response, a square root transformation for the BEC values was done. Variable selection using BIC based stepwise approach identified the following predictors as significant predictor variables at a 5% significance level (see Table 3a).

**Table 3a.** Regression model for Dataset2 consisting of influential conditions wherein weight was considered (age, vehicle, dose, time point, and weight).

|  | <u>Coefficient</u> | <u>Standard Error</u> | <u>t-value</u> | <u>p-value</u> |
| --- | --- | --- | --- | --- |
| <u>Intercept</u> | 13.43 | 1.54 | 8.74 | 0.00 |
| <u>Age</u> | 4.51 | 2.31 | 1.95 | 0.05 |
| <u>Vehicle</u> | -2.56 | 1.60 | -1.60 | 0.11 |
| <u>Dose</u> | 2.51 | 0.49 | 5.12 | 0.00 |
| <u>Time 90</u> | -2.75 | 0.64 | -4.29 | 0.00 |
| <u>Time 180</u> | -7.56 | 0.64 | -11.88 | 0.00 |
| <u>Weight</u> | -0.02 | 0.01 | -2.01 | 0.05 |
| <u>Age * Vehicle</u> | -7.28 | 2.39 | -3.04 | 0.00 |
| <u>Age * Time90</u> | 1.13 | 0.72 | 1.57 | 0.12 |
| <u>Age * Time180</u> | 2.61 | 0.72 | 3.62 | 0.00 |
| <u>Vehicle *</u><br><u>Weight</u> | 0.02 | 0.01 | 2.51 | 0.01 |
| <u>Dose * Time90</u> | 0.89 | 0.71 | 1.24 | 0.22 |
| <u>Dose * Time180</u> | 3.23 | 0.71 | 4.57 | 0.00 |

Thus, according to Table 3a, the final model is:

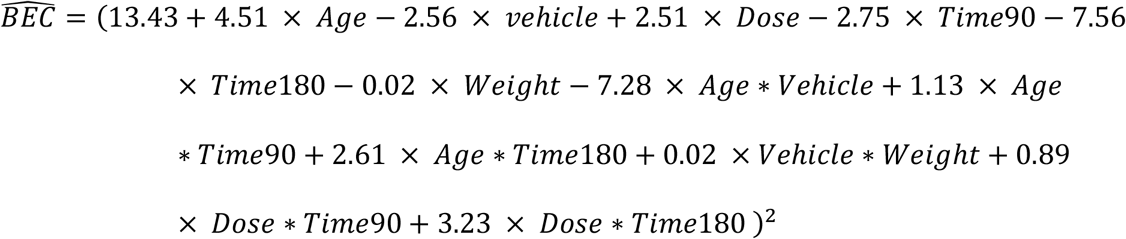

With this dataset, we had a relatively larger sample size of 203 and therefore, we also evaluated the stability of the model by cross-validation. A training-test set split was done 100 times by randomly assigning 80% of the observations into a training set and the remaining 20% of the observations into a test set. For each of the 100 training datasets created, we fitted the regression model as above and performed variable selection using a BIC based stepwise approach. The final regression model from each dataset was used to make a prediction on its respective test set and prediction accuracy was measured using R^2^. The overall mean R^2^ of the 100 datasets was 0.52 with a standard error of 0.1. We examined the top ten models with the highest R^2^ on the test set and they had a mean R^2^ of 0.66 with a standard error of 0.03. In 9 out of these 10 models, all the variables from the final model above were found to be significant while in the remaining model, the weight and vehicle interaction was not significant but the rest of the variables from the final model were significant. These data provide evidence for the robustness of the model.

##### Method 2

A second regression model was built using age, vehicle type, dose, and time point post-intubation as the predictor variables. Given that in the ANOVA conducted on CF estimation of BECs, weight was included in the calculation of BECs and eBECs (g EtOH/kg body weight), the regression analyses were conducted again with the weight variable excluded. Similar to the previous analysis indicator variables were created for categorical variables (age = 0 for adolescent and 1 for adult, vehicle = 0 for Boost^®^ and 1 for supersac, dose = 0 for 2 g/kg and 1 for 4g/kg, time_90 = 1 for 90 minutes and 0 for 45 and 180 minutes, and time_180 = 1 for 180 minutes and 0 for 45 and 90 minutes). We also examined two-way interactions among the predictor variables. Using a Box-Cox transformation on the response, a square root transformation for the BEC values was done. Variable selection using BIC based stepwise approach identified the following predictors as significant predictor variables at a 5% significance level (see Table 3b).

Thus, according to Table 3b, the final model is:

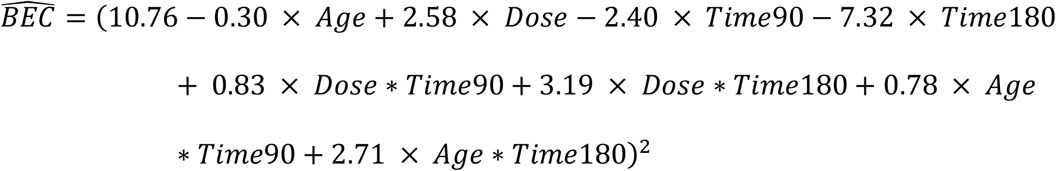

with an adjusted R^2^ of 0.63.

**Table 3b.** Regression model for Dataset2 consisting of influential conditions wherein weight was not considered (age, dose, time point).

|  | <u>Coefficient</u> | <u>Standard Error</u> | <u>t-value</u> | <u>p-value</u> |
| --- | --- | --- | --- | --- |
| <u>Intercept</u> | 10.76 | 0.43 | 24.76 | 0.00 |
| <u>Age</u> | -0.3014 | 0.50 | -0.60 | 0.55 |
| <u>Dose</u> | 2.58 | 0.50 | 5.14 | 0.00 |
| <u>Time_90</u> | -2.40 | 0.63 | -3.795 | 0.00 |
| <u>Time_180</u> | -7.32 | 0.63 | -11.61 | 0.00 |
| <u>Dose * Time90</u> | 0.83 | 0.73 | 1.13 | 0.26 |
| <u>Dose * Time180</u> | 3.19 | 0.72 | 4.41 | 0.00 |
| <u>Age * Time90</u> | 0.78 | 0.73 | 1.06 | 0.29 |
| <u>Age * Time180</u> | 2.72 | 0.72 | 3.76 | 0.00 |

Again, due to the relatively large sample size, stability of the model was examined through cross-validation using the same training-test set split approach as in the Method 1. The overall mean R^2^ of the 100 datasets was 0.51 with a standard error of 0.09. We examined the top ten models with the highest R^2^ on the test set and they had a mean R^2^ of 0.66 with a standard error of 0.05. In six out of these ten models, all the variables from the final model above were found to be significant while in the remaining model, the age and time interaction was not significant but the rest of the variables from the final model were significant. In three out of four models where the age and time interaction was not significant, the main effect of age became significant. These data provide evidence for reasonable robustness of the model.

### Dataset3 – CFs and estimated BECs after voluntary ingestion

#### BECs

The repeated measure ANOVA of the intake data collected on days 6 and 14 revealed no significant effects, whereas the ANOVA of the BEC data on these days revealed a main effect of day (F (1, 28) =4.498; p<0.05). BECs on day 14 (61.9 ± 6.2 mg/dl) were significantly lower than those on day 6 (84.2 ± 8.6 mg/dl) (See Supplemental Table 3).

#### CFs and eBECs

Given that this data set included intake and BEC samples on two days for each animal, a double repeated measure ANOVA was used to assess differences between BEC and eBEC across the two sampling days, with the variable comparing BEC to eBEC serving as a repeated measure nested within each sampling day. This ANOVA yielded a significant interaction of day X BEC/eBEC, suggesting one CF was not sufficient to accurately determine eBECs across days. When separate CFs were calculated for each day (day 6: CF = 20; day 14: CF = 14) and new eBECs calculated, the same double repeated measures ANOVA of these data revealed only a main effect of day (F (1, 31) =9.485; p<0.005), with BECs and eBECs being lower on day 14 than day 6, but BECs not differing significantly from eBECs on either day (P33: BEC = 84.2 ± 8.6 mg/dl; eBEC = 83.9 ± 5.1 mg/dl; P41: BEC = 61.9 ± 6.2 mg/dl; eBEC = 57.9 ± 3.1 mg/dl).

#### Results of regression analyses

Because this is a repeated measures study wherein BECs were analyzed at 2 time points within the same animals, an initial regression model was build using sex, tip-type, mean daily intake at P33 day and P41 day, and BEC values on the P33 day as the predictor variables and BEC values on the P41 day as the response variable. Indicator variables were created for categorical variables sex (sex = 0 for females and 1 for males) and tip-type (tip-type = 0 for ball-bearing tip and 1 for open-ended tip). Using a Box-Cox transformation on the response, a square root transformation for the BEC values on P41 day was done. Variable selection using a BIC based stepwise approach identified sex and mean daily intake on P41 day as a significant predictor variable for predicting BEC values on P41 day at a 5% significance level (see Table 4). Hence, according to Table 4, the final model is:

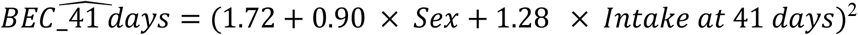

with an adjusted R-square of 0.73.

**Table 4.** Regression model for Dataset3 consisting of influential conditions such as Day 14 intake.

|  | <u>Coefficient</u> | <u>Standard Error</u> | <u>t-value</u> | <u>p-value</u> |
| --- | --- | --- | --- | --- |
| <u>Intercept</u> | 1.72 | 0.70 | 2.45 | 0.02 |
| <u>Sex</u> | 0.90 | 0.43 | 2.11 | 0.05 |
| <u>Intake at P41</u><br><u>day</u> | 1.28 | 0.17 | 7.76 | 3.11e-08 |

## Discussion

Three datasets were analyzed to determine factors important for deriving CFs for use in estimating BECs: subject age and sex, vehicle in which EtOH is administered, chronicity of alcohol exposure and post-exposure timing of blood collection. Surprisingly, our analyses using CF revealed no sex effects in either dataset where sex was investigated as a variable (Datasets 1 and 3). Likewise, in the CF analyses of age (see Datasets 1 and 2), this variable revealed few effects, with the only age difference observed evident late in the elimination phase (at 180 min. in Dataset2). Methodological variables including vehicle used and intake tube tip-type likewise were inconsequential upon CF outcomes. Despite the robustness of these eBEC calculations against variation in age and sex, eBECs were critically dependent on dose (Dataset2) and post-administration-to sample collection timing (Datasets 1 and 2). In an adolescent drinking sample (Dataset3), chronicity also affected accuracy of eBEC estimates via CF analysis, a factor that is rarely considered when deriving eBECs in either human or laboratory animals.

With regard to the regression analyses, Dataset1 only revealed an effect of time point. This finding was similar to the CF analysis. Dataset2 method 1 revealed influences of age, dose, weight, vehicle and time point whereas method 2 revealed findings more similar to the CF analysis, factoring in only age, dose and time point. The Dataset3 regression concluded that sex and P41 intake were important factors to consider when estimating BECs, a conclusion quite different from that reached in the CF analysis wherein sex did not play a critical role but P33 BECs did. Across 3 diverse datasets, we were able to derive estimates that can be used to approximate BECs when actual blood sample collection is not feasible, although the specific variables critical for deriving these estimates varied with the experimental conditions and, to some extent, across analysis approaches (CF vs. regression analyses).

Regarding age, BECs from blood samples collected after short time periods (around the time that BECs are approaching, plateauing, or declining from peak (e.g., 10, 30, or 45 minutes)) were little affected by age when comparing adolescents and adults; however, differences emerged between adolescents and adults when samples were collected 180 minutes post-challenge. The regression analyses confirmed that Dataset1 did not require a coefficient for age to determine eBECs at 10 and 30 minutes; however, this age distinction was important for determining eBECs in Dataset2 wherein longer time points, such as 180 min, were assessed. These data suggest that there are age-dependent differences in EtOH clearance/metabolism, with adolescents clearing EtOH from their system sooner than their adult counterparts. Importantly, adolescents and adults did not require separate CFs at the approximate time at which EtOH reaches its peak level within the blood. However, age may influence peak BECs at the same EtOH exposure level following chronic, or sub-chronic, exposure. A study that used repeated binge-level access to a nutritious EtOH diet found that, although adolescents consumed significantly more EtOH across all 4 days relative to adults, adolescents did not differ from adults in their peak BEC (Morris et al., 2010).

No sex differences emerged when calculating CFs for eBECs in Datasets 1 or 3, or in the regression analysis of Dataset1, although the regression analysis in Dataset3 did in fact use sex as an indicator for estimating BECs. Sex differences in EtOH clearance (Collins et al., 1975) and BECs (Seidl et al., 2000) have been reported in human and rodent studies, although these findings are not ubiquitous (e.g., (Goist Jr and Sutker, 1985, Kelly et al., 1987)). This may be due in part to higher availability of alcohol dehydrogenase in men than in women (Pastino and Conolly, 2000), and the reported more rapid absorption/diffusion rate from the stomach after i.g. EtOH in females than males (Robinson et al., 2002). Animal studies have shown that females consume greater quantities of EtOH per kilogram body weight (Lancaster et al., 1996, Lancaster and Spiegel, 1992). While males and females may differ in their time to regain consciousness from EtOH-induced sleep, their BECs at time of awakening do not differ, suggesting sex differences in EtOH metabolism (Collins et al., 1975). The difference between the CF analysis and the regression analysis confirm that there is not a simple relationship between sex and estimated BEC.

In the datasets examined, only Dataset3 addressed the contribution of chronic alcohol use on estimating BECs. Animals displayed a decrease in BEC relative to intake at P41 when compared to BECs and intakes at P33 in the same animals; therefore, a lower CF was necessary to estimate BECs at P41. There are two possibilities for these differences: age and weight at time of sample collection or amount of alcohol exposure. Because the only other age effect evident in these studies emerged late in the elimination phase (Dataset2), it seems most likely that the differences in eBECs from P33 to P41 reflect the emergence of metabolic tolerance. However, the regression analysis did not result in an indicator variable of P33 intake or BEC suggesting that early alcohol intake may not necessarily exert an influence on BECs observed later during the EtOH exposure period. It should be noted that several studies giving repeated i.p. injections of EtOH to adolescents and adult rats found adolescents to be relatively resistant to the emergence of metabolic tolerance relative to adults (e.g., (Broadwater et al., 2011, Varlinskaya and Spear, 2007)). Whether self-administration of EtOH accelerates expression of metabolic tolerance and does so differentially across age remains to be determined.

In Dataset2, regression analyses were conducted in two separate ways: one with weight included (method 1), the other without (method 2). An advantage of method 1 is precision; with information such as a specific animal body weight included in the indicator variables, eBECs may be more accurate but less generalizable. An advantage of method 2 is that weight is accounted for only once (in calculation of EtOH exposure in g/kg body weight). This model generated via method 2 was more general, and fit a wider range of variables. When weight was not included as a separate variable, the significant factors that influenced eBECs were age, dose and time point. When weight was included as a variable, an effect of vehicle emerged, suggesting that the absorption of EtOH in a sugary drink such as supersac versus a calorically dense chocolate drink like Boost^®^ may be different based on the weight of the subject. The vehicle was also affected by the age of the subject, as reflected by the age X vehicle intercept.

Although provision of calories with EtOH can delay absorption rate (Kalant, 1996), this is highly dependent on other factors including stomach content at the time of ingestion, a variable that was not controlled in the present study. Human studies have shown that EtOH absorption is more rapid with distilled spirits relative to beer when equal amounts of EtOH are provided (Mitchell Jr et al., 2014); however, this rate is opposite when food is present in the GI (gastrointestinal) tract (Roine et al., 1993). Rodent studies have found similar findings to Dataset2 in that, while sucrose increases the palatability of the EtOH solution and thus increases intake, provision of EtOH in a sucrose solution does not alter the ratio of intake to BECs 30 minutes after exposure (Czachowski et al., 1999). A limitation to this study was the early point of BEC collection, most likely during the absorption phase before peak BEC was achieved. Moreover, although EtOH was administered at the same time of day for all subjects in Dataset2, another limitation is that the presence of food in the GI tract was not controlled which could have obscured modest differences in EtOH pharmacokinetics between the two different vehicles. Finally, the weight X vehicle and age X vehicle interactions in the method 1 regression analysis did not reveal at which time point did weight mattered most and for which age group.

Because the datasets each used a different method of administrating EtOH and were conducted at different times, statistical comparisons across studies were not conducted. However, inspection of the summary data across these studies (see Table 1) demonstrates the notably higher CFs necessary with i.p. administration, reflecting the not surprisingly higher BECs and rapid absorption of EtOH from this route. CFs for Datasets 2 (i.g. intubation) and 3 (voluntary intake) were lower, demonstrating the slower uptake and/or more rapid metabolism through the “first-pass effect” associated with absorption through the GI tract. Interestingly, administration through the GI tract through oral self-administration versus i.g. intubation did not yield comparable CF’s, with the i.g. administrations of Dataset2 in general requiring about a two-fold higher CF than self-administered EtOH (Dataset3) to generate eBECs at roughly similar times post access (45 min. vs. ~30-35 min. from the start of drinking, respectively). These differences are likely due to differences associated with administering a bolus of EtOH quickly as opposed to voluntary consumption in which EtOH is administered more slowly over a period of time (up to 30 minutes in our dataset).

At first blush, deriving eBECs from CFs seems somewhat more complex in these rodent studies than the short list of variables considered in human studies where estimates are calculated from: time from the start of drinking episode, number of drinks consumed, gender, and weight (Hustad and Carey, 2005). These same variables were also found to be important when estimating BECs in rodent studies, with dose/exposure level and the timing of assessment exerting clear effects, body weight represented by always expressing exposures in g/kg, and a modest sex effect emerging in one of the regression analyses. However, interactions between dose, time post-challenge, and age (see Dataset2) complicated derivation of a single, simple formula to estimate BECs in these rodent studies, particularly late in the elimination phase. The variable of chronicity also was found to likely play a major role, a factor that has rarely been considered in human studies except in cases of liver damage where alcohol metabolism may be notably compromised (Shukla et al., 2008). A final important factor was route of administration (a variable irrelevant for human studies of oral self-administration), with this variable playing a major role when deriving CFs to estimate BECs in rats. Our formula is a first step into understanding how each of these factors plays a role in EtOH intoxication and metabolism. Moreover, many similarities between CF analyses and regression models suggest that there may be several ways to estimate BECs that may vary in their generalizability and accuracy.

Taken together, these datasets illustrate several important factors that influence BECs and the ability to accurately predict eBECs. Dose and chronicity are critical components that will likely influence eBECs and must be considered when estimating BECs while also taking into consideration the age, sex, and possibly other procedural variables (e.g., food/water deprivation). Another important facet of eBEC determination is the timing of sample collection mapped along the pharmacokinetic timeline (i.e. metabolism and elimination rates). Clearly, considerable caution is necessary when, in the absence of blood collection and analysis, BECs instead are roughly estimated by referring to BEC data reported in other studies at seemingly comparable EtOH exposure levels.

