## Supplemental Tables 1-3 for "Critical Variables to be Considered when Attempting to Estimate Blood Ethanol Concentrations in Rats from g/kg Exposure Data"

**Supplemental Table Legends**

**Supplemental Table 1.** Average BECs that were collected for Dataset1 for each age, sex, and time point group.

**Supplemental Table 2.** Average BECs that were collected for Dataset2 for each age, vehicle, dose, and time point group.

**Supplementary Table 3.** Average BECs that were collected for Dataset3 for males and females at P33 and P41 and their corresponding intakes.

**Supplementary Table 1.** Dataset1 Average BECs for each condition.

| Age | Sex | Time point | Average BEC |
| --- | --- | --- | --- |
| Adult | Male | 10 min | 119.8 ± 1.6 |
| Adult | Female | 10 min | 124.9 ± 6.7 |
| Adolescent | Male | 10 min | 114.8 ± 2.2 |
| Adolescent | Female | 10 min | 118.7 ± 2.3 |
| Adult | Male | 30 min | 102.0 ± 7.0 |
| Adult | Female | 30 min | 90.6 ± 7.1 |
| Adolescent | Male | 30 min | 96.3 ± 2.7 |
| Adolescent | Female | 30 min | 93.5 ± 4.3 |

**Supplementary Table 2.** Dataset2 Average BECs for each condition

| Age | Solution | Dose | Time point | Average BEC |
| --- | --- | --- | --- | --- |
| Adolescent | Boost® | 2 g/kg | 45 min | 105.0 ± 2.5 |
|  |  |  | 90 min | 71.8 ± 5.9 |
|  |  |  | 180 min | 12.1 ± 9.2 |
|  |  | 4 g/kg | 45 min | 163.6 ± 11.9 |
|  |  |  | 90 min | 140.0 ± 10.8 |
|  |  |  | 180 min | 87.2 ± 10.8 |
|  | Supersac | 2 g/kg | 45 min | 135.2 ± 13.4 |
|  |  |  | 90 min | 78.1 ± 7.0 |
|  |  |  | 180 min | 22.2 ± 9.3 |
|  |  | 4 g/kg | 45 min | 201.8 ± 27.3 |
|  |  |  | 90 min | 127.5 ± 8.8 |
|  |  |  | 180 min | 123.8 ± 28.1 |
| Adult | Boost® | 2 g/kg | 45 min | 108.6 ± 15.2 |
|  |  |  | 90 min | 87.7 ± 8.2 |
|  |  |  | 180 min | 49.7 ± 9.3 |
|  |  | 4 g/kg | 45 min | 173.6 ± 25.9 |
|  |  |  | 90 min | 166.2 ± 29.0 |
|  |  |  | 180 min | 114.8 ± 20.3 |
|  | Supersac | 2 g/kg | 45 min | 112.1 ± 8.7 |
|  |  |  | 90 min | 64.6 ± 8.8 |
|  |  |  | 180 min | 42.2 ± 5.7 |
|  |  | 4 g/kg | 45 min | 182.2 ± 21.9 |
|  |  |  | 90 min | 154.6 ± 8.6 |
|  |  |  | 180 min | 134.2 ± 11.8 |

**Supplementary Table 3.** Dataset3 Average intakes within each sex and P33 and P41 intakes and BECs

|  | Mean Intake Across 14 Days | P33 Intake (Day 6) | P33 BEC  (Day 6) | P41 Intake (Day 14) | P41 BEC  (Day 14) |
| --- | --- | --- | --- | --- | --- |
| Males | 4.2 ± 0.7 | 4.4 ± 0.5 | 81.0 ± 14.9 | 4.4 ± 0.3 | 71.9 ± 8.4 |
| Females | 3.5 ± 0.5 | 4.0 ± 0.2 | 87.3 ± 9.1 | 3.8 ± 0.3 | 51.9 ± 8.8 |
